# Metal Ions Guide the Production of Silkworm Fibers

**DOI:** 10.1101/2023.08.10.552775

**Authors:** Ori Brookstein, Eyal Shimoni, Dror Eliaz, Ifat Kaplan-Ashiri, Itay Carmel, Ulyana Shimanovich

**Affiliations:** Department of Molecular Chemistry and Materials Science, Faculty of Chemistry, Weizmann Institute of Science, 7610001 Rehovot, Israel; Department of Chemical Research Support, Faculty of Chemistry, Weizmann Institute of Science, 7610001 Rehovot, Israel; Department of Chemical and Structural Biology, Faculty of Chemistry, Weizmann Institute of Science, 7610001 Rehovot, Israel

**Keywords:** Silk, Metal ions, Spinning, Rheology, Mechanical properties

## Abstract

Silk fibers’ unique mechanical properties have made them a desirable material for various applications, from medical to optical materials and even in sensing. Yet, to date, no synthetic method has come close to reproducing this remarkably strong biomaterial due to the complexity and insufficient understanding of the mechanism of silk fiber formation. While ions are known to play a key role in the production of natural silk fiber, how they do so has thus far eluded discovery. Here we report that a broad composition of metal ions guides structural transformations in the silk fibroin protein inside the silkworm silk gland. By using a combination of cryo-electron microscopy techniques coupled with elemental analysis, we followed the changes in the composition and spatial localization of metal ions inside the silk gland. We observed that ions are homogenously dispersed during the initial stages of silk secretion and storage inside the silk gland, but once the fibers are spun, the ions delocalize from the silk fibroin fiber core to the sericin coating gum layer. This shift in ion localization is accompanied by the alignment of protein chains and an increase in silk feedstock viscosity inside the silk gland - changes that make the protein more sensitive to shear and enable the initiation of the liquid-to-solid transition in the silk. Moreover, the selective doping of the spun silk fibers with metal ions modifies their mechanical performance. These findings highlight the importance and the dynamic role of metal ions in the evolution of silk fibers’ mechanical properties, enhance our understanding of the mechanism of silk fiber formation, and lay the foundations for developing new concepts in biomaterial design.

## Introduction

Silk fibers are known for their unique combination of mechanical characteristics of strength, extensibility, and toughness. These features result from silk’s highly hierarchical structure, shaped by the protein’s self-assembly during the natural fiber’s spinning. The entire production of spider and silkworm (e.g., *B. mori*) silk fibers occurs inside the silk gland, a long tube-shaped organ made of epithelial cells that envelope an inner lumen where the liquid silk protein (fibroin in the case of *B. mori* silk) feedstock flows^1,2^. Fibroin is first synthesized in the cells and secreted to the lumen in the posterior section of the gland. The liquid protein then flows to the middle section, where it is stored at conditions that prevent unwanted premature fibrillation. Upon the silk feedstock flow reaching the anterior part of the silk gland, the soluble protein transitions into a highly-organized β-sheet-rich solid fiber^1,3–5^. The protein storage, structural transition and fibrillation stages are all governed by the fluid rheology and the effects of shear, elongational flow, pH, and metal ions^1,6–12^.

While the impact of most of these factors has been extensively studied^3,6,8,13–16^, the role of metal ions in this process remains largely unclear. Of the studies concerning the metal ions, most have focused on the effect of their interaction with fibroin on the secondary structure of the soluble protein. In silkworms, metal ions such as Na, Mg, Zn, Cu, and Fe have been suggested to induce a conformational transition of reconstituted silk fibroin (RSF) from a random coil to a β-sheet structure^17–21^. However, it has been reported that while a low concentration of Ca ions permits β-sheet formation in RSF, a higher concentration inhibits it^22,23^. Other studies have demonstrated how the presence of metal ions influences silk fiber’s secondary structure and mechanical properties^24–26^. On the one hand, Wang et al.^25^ have shown that genetic manipulation to suppress the amount of K and Na in silkworm’s silk feedstock increases the random coil conformation and lowers the Young’s modulus and toughness of the spun fibers. On the other hand, reducing the Ca content raises the number of α-helix and β-sheet structures in silk fibers and improves their strain-to-failure and toughness.

More recently, a study relating silk feedstock flow properties with metal ions has suggested that a high concentration of the bivalent ion Ca promotes suitable storage conditions for fibroin by creating ion bridges, increasing feedstock viscosity, and inducing gelation. Yet, it appears that other ions in the silk gland, such as the monovalent K and, to a lesser extent, Na and Li, set the conditions for spinning by breaking down ion bridges and lowering the feedstock viscosity. This lets the proteins flow, increases shear sensitivity, align and form β-sheet conformations^10,12,21^.

Several studies of the natural silk-spinning process *in vivo* have indicated that metal ions are regularly present in the silk gland, and that their concentration changes along the duct^21,27–29^. Zhou et al. demonstrated that Na, Mg, Zn, and Cu concentrations increase progressively from the posterior-middle to the anterior-middle sections of the silk gland, while that of Ca decreases^21^. This consistent trend in ions’ concentration suggests that they are not merely contaminations but essential factors in the silk-spinning process. However, insight into the actual distribution of metal ions in the silk materials and during their processing is needed to understand how they participate in this natural spinning process.

Here, we used an array of high-resolution analytical, spectroscopy, and microscopy techniques to comprehensively study metal ions’ content and spatial distribution inside the silk gland and silk fiber. We then examined their impact on silk feedstock characteristics and fiber’s physical properties. Our investigation includes an analysis of the various sections of the silk gland, relevant to different stages of the silk protein’s spinning. The analysis was performed to best preserve the native states of the materials and biological tissues and to separate the different regions of interest. The metal ions’ total concentration and spatial localization were found to change along and across the silk gland (**Figure 1a**). The ions are homogeneously distributed in the posterior tissue (**Figure 1a**), but gradually increase in concentration from the posterior-middle to the anterior-middle sections of the silk gland. While flowing from the anterior-middle to the anterior sections, the metal ions relocate to the cuticular intima (sericin) and epithelial cells (**Figure 1c**). In the final product, naturally spun silk fibers, most metal ions are localized in the sericin coating layer and are much less abundant in the fibroin core (**Figure 1d**). This observation suggests that both factors; 1) the change in pH that might midify the protonantion state of the silk protein, and 2) the presence of sericin component that can act as an ion sink, change ions’ localization at the silk spinning stage. These variations in ion localization inside the gland were reflected by changes in silk feedstock viscosity, enabling to switch between a shear-insensitive storage state to a shear-sensitive spinning mode, namely, the liquid-to-solid transition in the silk. Finally, the selective doping of the spun silk fibers with metal ions changed the fibers’ mechanical characteristics.

**Figure 1.**
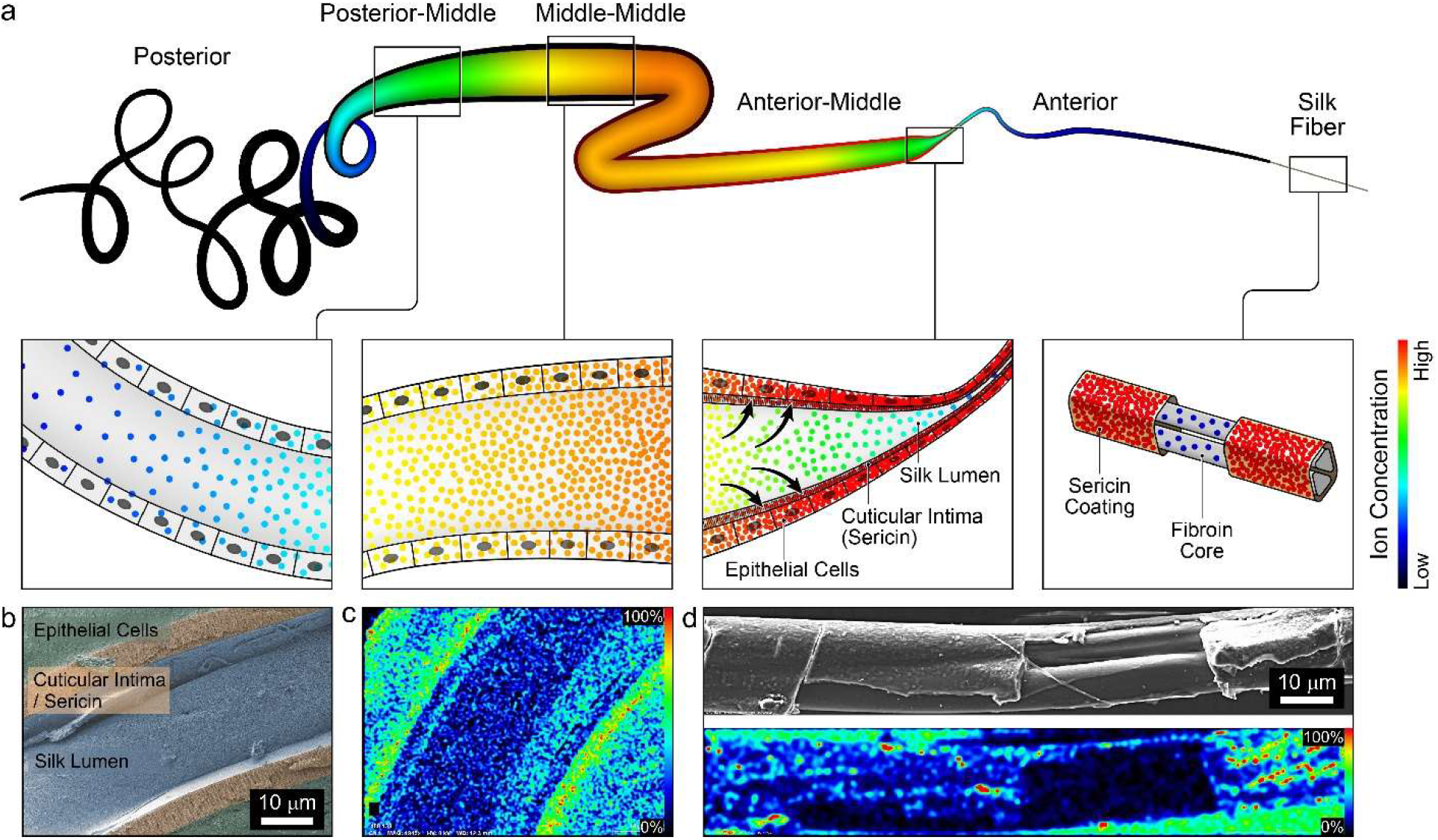
Metal ions’ distribution during silk processing. **(a)** Schematic illustration of the change in the overall metal ion distribution through the silk gland and in the silk fibers. Colors reflect the level of concentration, as indicated in the color scale to right. **(b)** Freeze-fractured cryo-SEM image of silkworm’s anterior section of the silk gland, showing its three components: epithelial cell layer, cuticular intima and silk lumen; false colors. **(c)** General example of a cryo-SEM-EDS normalized intensity map of the total ions’ distribution inside the anterior silk gland. **(d)** SEM image of a silk fiber with an exposed region of the core fibroin fibers (top) and an EDS heat-map image of the overall metal ion distribution in this fiber (bottom); same color scale as in **(a)**.

**Table 1:**
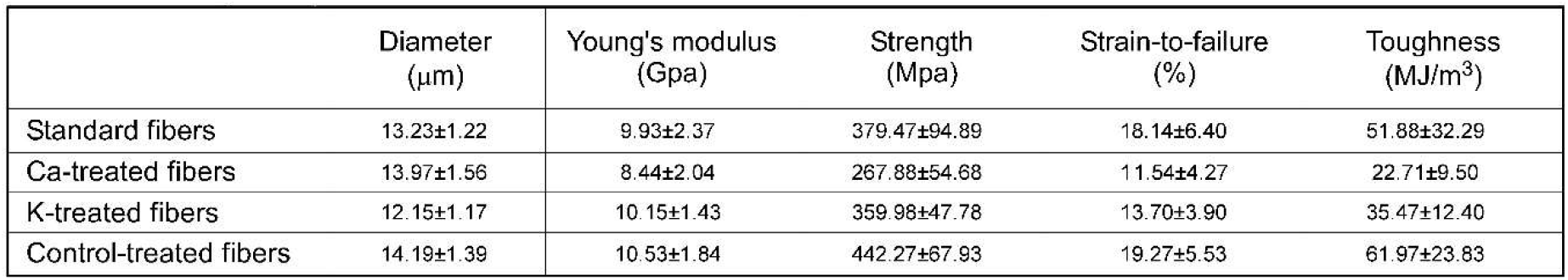
Summary of tensile measurements of silk fibers with metal ions treatments.

## Materials and methods

### Preparation of degummed and reconstituted silk

5 g of *Bombyx mori* silk cocoons were cut into small pieces and soaked twice, for 15 min each time, in 2 liters of boiling Milli-Q water with 0.02 M sodium carbonate. Next, the degummed fibers were washed exhaustively in Milli-Q water and then left to dry for three days at room temperature.

A reconstituted silk fibroin (RSF) solution was prepared by dissolving dried fibers in 9.3 M LiBr (20% w/w) at 65°C for 4 hours. The solution was dialyzed in a 10 kDa dialysis bag placed in distilled water for 72 hours while changing the water three times a day. The solution was then centrifuged twice at 12,000x g at 4°C for 20 min, with the precipitates removed after each cycle, and then the solution was stored at 4°C. The RSF concentration was calculated by weighing a known volume after drying and measuring the protein solution’s light absorption using a nano-drop instrument.

### Extraction of silk glands and silk feedstock

*Bombyx mori* larvae at their fifth instar were anesthetized with N_2_ for 20 min and then rapidly dissected by removing their head and applying a longitudinal dorsal incision. The silk glands were gently extracted and rinsed with Mili-Q water. For elemental analysis, each gland was briefly cut into five sections: posterior (P), posterior-middle (PM), middle-middle (MM), anterior-middle (AM) and anterior (A). The silk feedstock was extracted by placing each gland section in Mili-Q water, to allow the viscous solution to flow out, and then gently peeling off the epithelial layer. For the cryo-SEM (scanning electron microscope) imaging, the anterior section (A) was briefly removed and frozen, as described below.

### Metal ion quantification using inductively coupled plasma mass spectrometry (ICP-MS)

The content of metals in silk fibers and silk glands or reconstituted silk fibroin was measured using an ICP-MS analysis. The samples were dried in a 100°C oven for three days, weighed, and then ashed in a 450°C furnace for an additional three days. Each sample was dissolved in 3 ml of 3% nitric acid overnight, sonicated for 60 min, and filtered through a 0.22 μm PVDF filter before the ICP-MS analysis (7700 Series, Agilent Technologies, Santa Clara, CA, USA). The system was calibrated with 0, 5, 10, 20, 50, 100, 500, and 1000 ppb certified ICP-MS standard calibration solutions (IV-STOCK-21, Inorganic ventures, USA) prepared in 3% (v/v) nitric acid (≥99.999% trace metals basis, Sigma-Aldrich). An internal standard solution (3% (v/v) HNO_3_) containing 1000 ppb of Gadolinium was injected throughout the measurements. The elements were determined using an Octapole Reaction cell (ORS3). The average results of each element’s concentration were calculated for triplicate or duplicate measurements using MassHunter 4.1 Workstation Software for 7700 ICP-MS (Agilent Technologies Inc. 2015).

The concentration of each metal ion was calculated by dividing its measured weight in the sample (where the measured ppb was divided by the sample’s volume) by the total weight of the sample. The number of each ion type per protein chain was calculated by dividing its number in the sample (calculated using its total measured weight and its molecular weight) by the number of fibroin protein monomers in the sample (calculated using the total sample weight and the molecular weight of fibroin (415 kDa), based on the protein’s light and heavy chains’ amino acid sequence from https://www.uniprot.org/uniprot/P05790, https://www.uniprot.org/uniprot/P21828).

### Silk fiber imaging and elemental analysis by SEM-EDS

Silk fibers were mounted on aluminum or titanium stubs using a two-sided carbon tape and then coated with 6-nm-thick carbon using a Safematic CCU-010 HV coater. High-resolution SEM images were acquired using a Zeiss Ultra 55 SEM. Following the imaging, energy-dispersive X-ray spectroscopy (EDS) hypermaps were acquired with a four-quadrant retractable detector (Bruker QUANTAX FlatQUAD EDS) mounted above the sample. Imaging and EDS acquisition were obtained at acceleration voltages of 2, 4, or 8 kV (60 μm aperture), sufficient for generating a characteristic X-ray for the elements of interest. The normalized X-ray intensity maps of the element lines C-Ka, O-Ka, S-Ka, P-Ka, and Ca-Ka were produced from the hypermap data.

### Silk gland sample preparation for electron microscopy High-pressure freezing of silk gland

Silkworm anterior gland samples were cut using a razor blade and placed on aluminum discs (Wohlwend Engineering Office, Switzerland). For freeze-fracture, samples were sandwiched between two aluminum discs 100 µm in depth, to fit the diameter of the gland sample. The discs’ surface was scratched with a razor blade in order to prevent separation between the discs and the sample during the freeze-fracture process. To prevent air bubbles, empty spaces between the sample and the disc walls were filled with hexadecane or Mili-Q water. For freeze-substitution, samples were placed on a disc 200 µm in-depth and covered with a flat disc. The sandwiched sample was frozen in an EM ICE high-pressure freezing (HPF) machine (Leica Microsystems, Vienna, Austria) and then kept in liquid nitrogen for future processing.

### Freeze fracture cryo-SEM imaging and EDS analysis of silk glands

The high-pressure frozen samples were mounted on a holder and transferred to a BAF 60 freeze-fracture device (Leica Microsystems, Vienna, Austria) using a VCT 100 Vacuum Cryo Transfer device (Leica). After fracturing at a temperature of −120°C, samples were transferred to an Ultra 55 SEM (Zeiss, Germany) using a VCT 100 cryo shuttle, and observed using a secondary electron in-lens detector at an acceleration voltage of between 1–2 kV at a temperature of −120°C. Some structures could only be revealed after sublimation (etching) at (−90)–(−80)°C for 10–20 min. Elemental analysis was done, as described above, using a FlatQuad EDS detector at acceleration voltages of 2, 4, 8, or 15 kV, sufficient for generating a characteristic X-ray spectrum for the elements of interest.

### Rheological measurements of metal-doped RSF solutions

RSF solutions (final protein concentration of 2%) were placed in dialysis tubes and dialyzed against 0.5 L of KCl, CaCl_2_, or NaCl solutions at different concentrations to reach protein-to-metal molecular ratios of 1:3, 1:30 and 1:300. To measure the number of ions binding to fibroin, samples doped with high ion concentrations were dialyzed against Mili-Q water; the water was changed three times to remove any non-binding ions. The samples were then measured using ICP-MS, as described above.

Rheological measurements were done using an HR-20 Discovery Hybrid Rheometer (TA Instruments, US) with a 40 mm diameter aluminum parallel plate geometry at 25°C. Samples of 0.5% RSF were placed on the instrument’s surface, and the geometry was lowered at the slowest speed possible to a gap of 100 µm, to avoid applying high shear to the sample. A small amount of distilled water was placed around the sample and covered with a loose-fitting solvent trap to minimize water loss and prevent skin formation. The procedure started with a constant shear rate of 1 s^-1^ for 100 seconds to evenly distribute the liquid and avoid residual stresses in the sample. The initial “pre-shear” viscosity was calculated by averaging the measurement’s last 30 seconds.

Next, an oscillatory frequency sweep was done from 100 rad/s to 0.1 rad/s at a shear strain of 0.02 (within the material’s linear elastic region). The dynamic viscosity of the sample was then measured during two repeated flow sweep tests of a gradual increase and decrease in shear rate from 0.1 to 500 s^-1^, recording six points per decade with an average time of 10 seconds per point. The sample was then subjected to a second constant shear rate of 1 s^-1^ for 100 seconds. The “post-shear” and “relaxed” viscosity was calculated by averaging the measurement’s last 30 seconds. Finally, a second oscillatory frequency sweep was done with the same parameters as before. Data were analyzed by 2-way ANOVA followed by a Tukey HSD multiple comparisons test.

### Mechanical measurements of metal-doped silk fibers

Silk fibers were straightly placed on paper frames (gauge length: 10 mm) using double-sided tape and glued with epoxy. The diameter of each tested fiber was measured by automatically imaging each sample along its entire length with a light microscope occupied with a mechanically controlled stage. The local thickness was measured in each image, and the average diameter along the whole length of the sample was calculated using a homemade Fiji^30^ macro. The macro first segment the fiber into a binary mask by applying background subtraction using rolling ball, followed by segmentation with auto-threshold (Li method). Local thickness^31^ is calculated for the binary mask and measured along the center line of the fiber, which is extracted from a cleaned skeleton of the binary mask. We erode the mask prior to skeletonization and discard skeleton regions at the edges of the fiber and those that are perpendicular to the fiber main orientation.

Tensile tests were conducted with an Instron 5965 universal testing system (UK) equipped with a 10 N load cell. The paper frame was held on the instrument with a pair of clamps, and the tensile stress was measured at a strain rate of 1 mm/min.

Data were analyzed by 2-way ANOVA followed by a Tukey HSD multiple comparisons test.

## Results and discussion

### Composition and spatial distribution of metal ions in silk fibers

We initially assessed the overall metal ion composition (i.e., the total number of metal ion elements) and localization in spun silk fiber using inductive coupled plasma mass spectroscopy (ICP-MS) and scanning electron microscope (SEM) coupled with an energy-dispersed X-ray spectroscopy detector (SEM-EDS). While ICP-MS is an elemental analysis technique that enables the detection and quantification of a number of different elements present in the sample, the SEM-EDS technique provides complementary information about the spatial localization of the elements inside the sample. Our ICP-MS analysis revealed that natural silk fibers (fibers composed of a fibroin core and sericin coating) contain a broad range of different elements. The most abundant ones are K, Ca, Na, and Mg (**Figures 2a and b**), a finding in agreement with previous reports^12,21,27^. Interestingly, we observed that the spatial distribution of the above-listed abundant elements along the sericin-coated fiber (**Figures 2c and supplementary Figure S1**) is not uniform, with element-rich and element-poor regions (**Figure 2d**). The relatively high amounts of the K and Ca elements compared to other elements has been explained by their ability to modify silk protein’s conformation, thus preparing the silk feedstock solution for spinning^19,22,23^. K is generally associated with the unwinding of the protein chains, which leads to the adopting a β-sheet conformation, associated with further fibrillation. Ca inhibits the fibrillar self-assembly process by keeping the protein molecules in a random-coil conformation. Our SEM-EDS analysis found that K and Ca tend to co-localize in the spun fibers, as opposed to Mg, which accumulates in distinct areas (**Figure 2d**). This may be indicative of the “active” protein-modifying role of K and Ca and as opposed to the less pronounced effects of other elements, including Mg, on the protein’s conformation. In addition to the presence of abundant elements (i.e., K, Ca, Na, and Mg), Fe, Zn, and Cu were also found in the sericin-coating layer of silk fiber. These elements appeared at much lower concentrations, even though they were previously suggested to affect silk protein’s secondary structure^20,21,27,32,33^.

**Figure 2.**
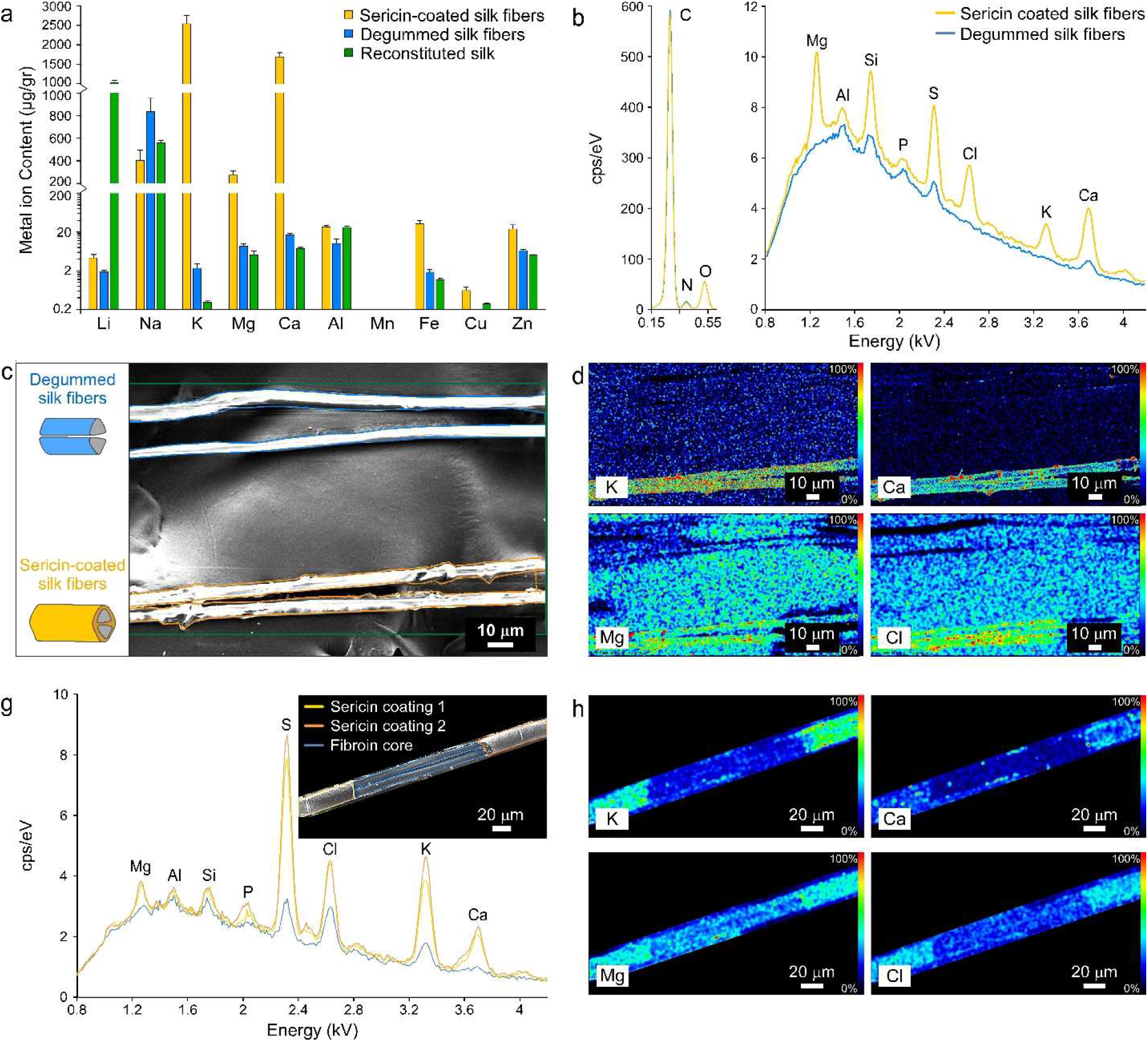
Metal ions in native and processed silk fibers. ***(a)*** *ICP-MS quantification of metal ions (n=6) and **(b)** the characteristic X-ray spectrum of elements obtained by SEM-EDS in silk fibers before (“Sericin-coated”; orange/yellow) and after (“Degummed”; blue) removing the native sericin coating, and after reconstitution (“Reconstituted”; green).**(c)** SEM-EDS image and the regions of interest in the silk fibers same colors as in **(a)**; scale 10 µm. **(d)** Normalized SEM-EDS intensity maps of the K, Ca, Mg and Cl elements’ spatial distribution in the silk fibers (same image as in **(c)**); color scale (0–100%) on the right of each image. Scale 10 µm. **(g)** Characteristic X-ray spectrum of elements in the different regions of silk fiber with exposed fibroin core, shown in the insert SEM image. **(h)** Normalized SEM-EDS intensity maps of the K, Ca, Mg, and Cl elements’ spatial distribution in the fiber (same image as the insert in **(g)**); color scale (0–100%) on the right of each image. Scale 20 µm. The EDS was acquired at an accelerating voltage of 2 keV for low energy elements (C, N O) and at an accelerating voltage of 8 keV for higher energy elements*.

Next, we compared the elemental composition in silk fibers that contain a sericin-coating layer versus degummed silk fibroin fibers lacking the sericin coating (**Figure 2c)**. We detected a striking difference between these two types of fibers: While the composition of the three most common elements in systems of organic nature, i.e., C, N, and O, in both fiber types was similar (**Figure 2b**), the concentration of the metal ions (weight of a given ion divided by the total weight of the material) was much higher in sericin-coated fibers compared to uncoated fibroin fibers (**Figures 2a, b**, and **supplementary Figure S1a**).

The standard degumming procedure, namely, removing the sericin coating layer, requires boiling silk fibers in an aqueous detergent-containing solution. Thus, to understand whether the reduction in the metal ions’ concentration in the degummed fibers is derived from the degumming procedure itself, we examined “sericin-defected” areas in native silk fibers, namely, regions where the sericin gum was either scratched or detached. These surface defects expose the fibroin core without changing the natural elemental composition of the entire fiber (**Figures 2g insert and supplementary Figures S2a** and **b**). The results from the SEM-EDS analysis of the sericin-defected fibers are consistent with the degummed fibers and show a distinct difference in the metal ions’ distribution: the sericin layer contained much higher concentrations of all the elements compared to the fibroin core (**Figures 2g, h**, and **supplementary Figure S2c)**.

Furthermore, our findings show that the degumming procedure, in which the sericin coating is removed, reduces the overall concentration of metal ions in the fibroin fibers (**Figure 2g**). This might potentially affect the fiber’s mechanical properties (see detailed discussion in the “Metal ions modulate the mechanical properties of silk fibers” section).

### The metal ion content in the silkworm silk gland

To further understand the rationale for the observed variations in metal ion distributions in the spun fibers and dynamics of such changes during the process of silk fiber formation, we performed an elemental analysis on the silkworm’s silk gland. This entailed dissecting and removing silk glands from adult silkworms (see Materials and Methods) and separating silk glands into their main sections: posterior, posterior-middle, middle-middle, and anterior-middle (**Figure 1a**). The tissue of each section was taken as a whole (namely, containing the silk lumen (fibroin-rich section), sericin cuticula intima and epithelial cells of the silk gland, as illustrated in **Figure 1b**) and measured to determine the concentration of the various elements by ICP-MS. This analysis revealed a similar elemental composition to that in spun silk fibers, with the most abundantly detected metals being K, Na, Mg, and Ca (**Figures 3a and b**).

**Figure 3.**
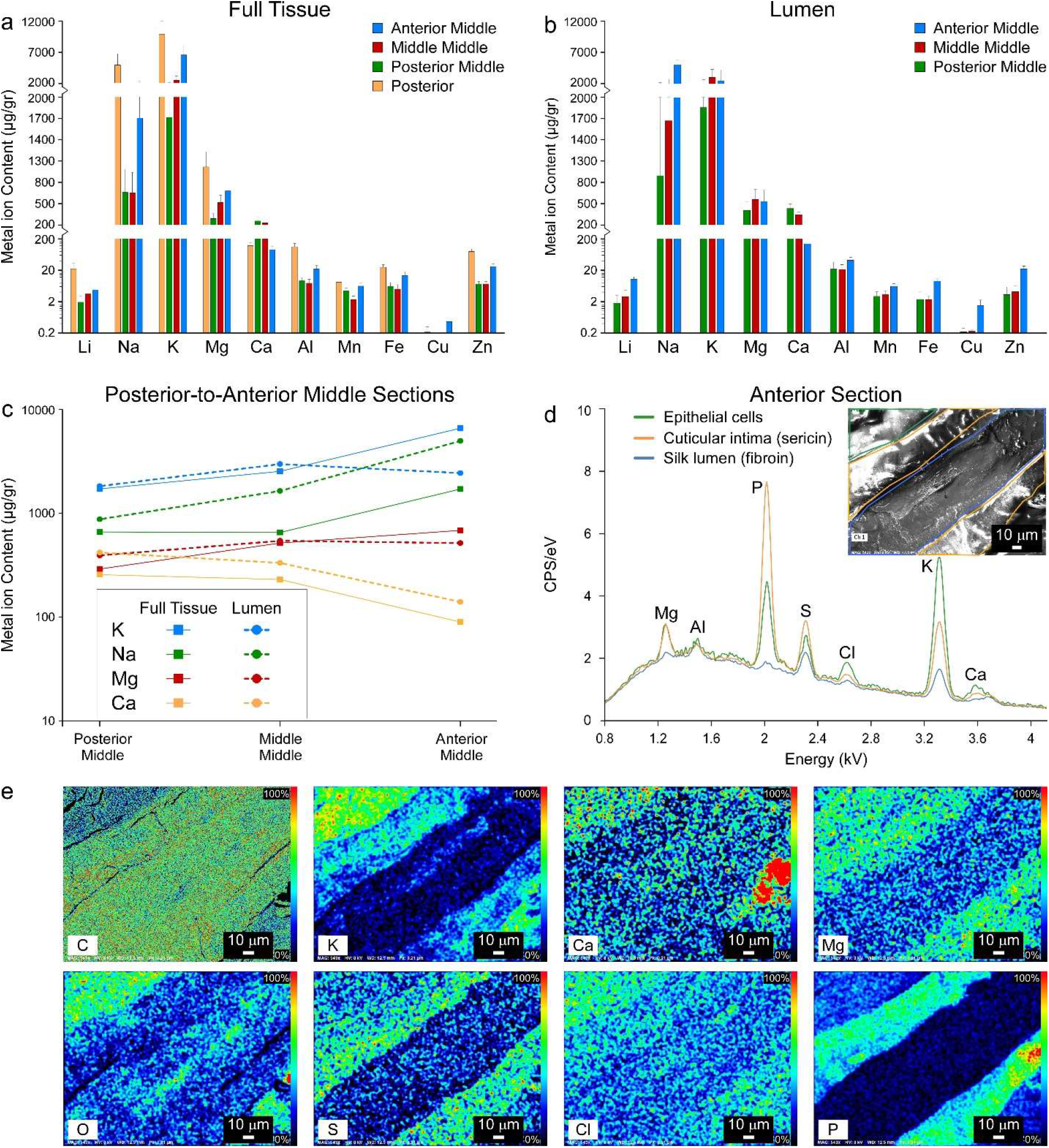
Concentration and distribution of elements along the silk spinning process. (a-b) ICP-MS quantification of metal ions content in the full gland’s tissue (a) and only inside the gland’s lumen (b) at the posterior section (yellow), posterior-middle (green), middle-middle (red) and anterior-middle (blue) sections. (c) The content of K, Na, Mg and Ca elements in the full gland’s tissue and only in the inner lumen. (d) Characteristic X-ray spectrum for elements of the anterior silk gland in the ephithelial cells (green), cuticular intima (yellow) and silk lumen (blue) regions, as shown in the inset cryo-SEM image. (e) Normalized cryo-SEM-EDS intensity maps of the elements’ distribution in the gland (same image as the insert in (d)); color scale (0–100%) on the right of each image. Scale 10 µm. The EDS was obtained at an accelerating voltage of 8 kV.

We further observed that the average metal ion concentration, throughout the entire gland tissue, is highest in the posterior section (**Figure 3a**). In this section, the gland’s epithelial cells synthesize the fibroin proteins and secrete them into the lumen. Therefore, we hypothesize that this high concentration of ions is related to fibroin’s secretion into the lumen and the stabilization (via preventing premature fibrillation) required for protein storage.

The ions’ concentrations sharply decrease from the narrow posterior section to the wider posterior-middle section of the silk gland. Surprisingly, this stark decrease is followed by a gradual increase from the posterior-middle to the middle-middle and all the way to the anterior-middle section of the silk gland (**Figure 3a**). A notable exception is the concentration of the Ca ions, which has the opposite trend, namely, lower in the posterior and the anterior-middle and higher in the posterior-middle and middle-middle sections. Previous *in-vitro* studies have suggested that changes in metal ions’ concentrations can be attributed to their ability to induce a structural transition in the silk protein ^10,12,21^. Thus, ions such as K, Mg, Zn, and Cu induce β-sheet formation. The increase in their concentration in the gland’s anterior-middle part can be associated with the random-coil-to-β-sheet transition and in preparation for the protein’s spinning. In contrast, the high Ca concentration can be associated with stabilizing the random-coil conformation connected with inhibiting fibrillation, which should make this ion critical for storing silk feedstock in the middle and posterior regions of the gland^21,23^.

To further relate the role of metal ions to silk protein processing, we compared their concentrations in the whole gland tissue and silk feedstock (i.e., the aqueous silk proteins in the fluid extracted from the silk gland’s lumen; **Figures 1a, b**, and **Figure 3a**). For this purpose, silk feedstock fluid was gently extracted from the different sections of silk glands of dissected silkworms (see Materials and Methods) and analyzed by ICP-MS. We observed no pronounced differences between the localization and concentration of most elements in the gland’s lumen versus epithelial cells along most of the silk gland, specifically from the posterior to the middle-middle sections; they were homogeneously distributed between the tissue and the feedstock. Changes in ion localization and concentration start to appear from the anterior-middle section onward (**Figures 3b and c**). For example, K and Mg ions’ concentrations at the anterior-middle section increased in epithelial cells but decreased in the feedstock (**Figure 3c**). This change indicates that metal ions start moving out of the lumen’s feedstock as it enters the anterior part of the silk gland. We believe that the difference between the ion concentration in the fibroin feedstock and in the rest of the anterior tissue is even more pronounced. As we have detected, there is a slight sericin “contamination” in the extracted fibroin liquid (**supplementary Figure S3**), which would increase the total amount of metal ions in the sample.

In the anterior section of the silk gland, which is closer to the spinneret, the amount of the protein is small, making the differentiation between the lumen and tissue parts technically impossible. We, therefore, measured the level of metal ions in this region using cryo-SEM-EDS (**Figure 3d**). To best preserve the sample in its native state, the anterior section tube was briefly cut and subjected to high-pressure freezing, followed by freeze-fracture and imaging by a cryo-SEM system coupled to a highly-sensitive EDS detector. We found a distinct uneven distribution of metal ions inside the anterior section (**Figures 3d and e**): the concentration was much higher in the cuticular intima (sericin) and the epithelial cells and significantly lower in the fibroin lumen. Thus, the ion concentrations in the lumen decrease as the silk feedstock flows from the anterior-middle to the anterior section. Previously, Wang et al.^29^ reported that ionic pumps operate in the gland’s anterior region, suggesting that it actively transports Na, K, and Ca across cell membranes. The distribution of ions observed in our analysis (**Figure 3e**) supports this hypothesis, as it indicates that ions flow from the lumen to the epithelial cells, thereby lowering the concentration in the silk feedstock. Our findings may also support the conjecture of Knight and Vollrath^28^ that ion reabsorption by epithelial cells in the spider gland relates to water uptake from the lumen, which is essential for the final stage of the fiber’s spinning process, accompanied by dehydration events.

The differences in the concentrations of ions between the lumen and the cuticular intima resemble those between the spun fiber’s fibroin core and sericin coating. The uneven distribution between those liquid phases highlights a difference in metal ions’ partitioning between the fibroin and sericin phases, which is an attractive subject for future study.

As the silk feedstock flows into the anterior part of the gland, it shifts from a “stable storage” regime to an “active spinning” state. This shift is dictated by the surrounding environmental conditions and applied forces (shear under elongational flow created during the spinning process). Thus, our results show that part of these conditions involves a deliberate reduction in metal ion concentrations. Nevertheless, the storage conditions for preventing premature fibrillation feature higher metal ion concentrations.

Interestingly, we observed that the metal distribution in the silk feedstock changed when fibrillation and bundle formation were induced during the gland’s dissections (**supplementary Figures S4 and S5**). Elements like K, Cl, and Ca were detected at higher concentrations in the fiber-like phase, which points to the role of those elements in the promotion or stabilization of silk fibrillation under shear.

### Silk Feedstock Rheology Modulated by Metal Ions

To obtain a perfectly oriented and hierarchical structure, the transition of silk from a liquid to a solid state must be gradual and highly controlled. Thus, the fluid characteristics are carefully tuned and controlled at each stage of the spinning process. To understand how the ionic environment modifies silk fluid characteristics in preparation for the spinning of the protein-rich liquid, we performed a set of *in-vitro* experiments.

First, we evaluated the most abundant metal ions’ binding propensity to fibroin (**Figure 4a**). Although the metal ions’ interactions with fibroin and their effect on its conformation have been previously studied^18,19,21,34–36^, the number of ions bound to each protein chain has not been determined. For this purpose, we doped RSF solution (a fibroin solution with a known and controllable ion composition) with high concentrations of K, Na or Ca ions (see Materials and Methods), evaluated the binding of the dopant ions to silk protein, and then tested their effect on silk fluid characteristics. The doped solutions were then washed with DDW (via dialysis) and analyzed by ICP-MS (**Figure 4a** and **supplementary Figure S6a**) to quantify the protein-bound ions.

**Figure 4.**
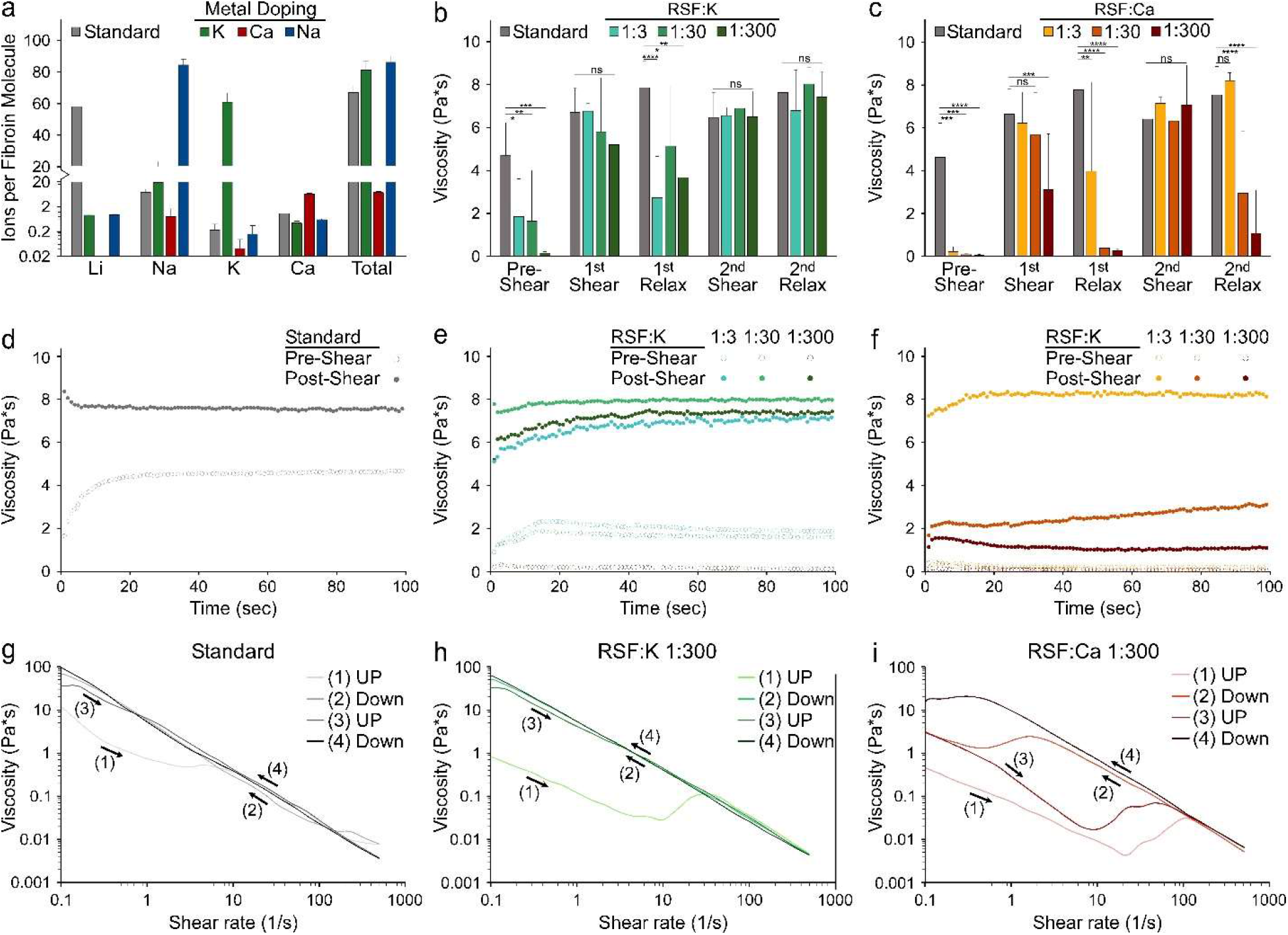
Modulation of silk feedstock properties by metal ions. ***(a)*** *Ratio of binding metal ions per fibroin molecule in reconstituted silk fibroin (RSF) solutions after metal-doping with Na, K, or Ca, measured using ICP-MS. **(b-i)** Shear viscosity measurements for the standard and metal-doped RSF solutions with fibroin-to-ion molecular ratios of 1:3, 1:30 and 1:300. Averaged viscosity at 1s^-^*^1^ *shear rate of standard RSF and RSF doped with K **(b)** or Ca **(c)**, before, immediately after or after a relaxation period following two cycles of recurring high shear. **(d-f)** The change in viscosity over time at 1s^-^*^1^ *shear rate before and after two cycles of high shear for the standard **(d)**, K-doped **(e)** or Ca-doped **(f)** RSF. **(g-i)** The shear viscosity plotted against the shear rate from 0.1 s^-^*^1^ *to 500 s^-^*^1^ *on two cycles of increasing and then decreasing rates, for the standard RSF **(g)**, 1:300 K-doped **(h)** and 1:300 Ca-doped **(i)** RSF solutions. ns (not significant) - P>0.05, *-P<0.05, **-P<0.01, ***-P<0.001, ****-P<0.0001*.

As a reference, we used an aqueous non-doped RSF solution. The ICP-MS analysis of non-doped RSF (“Standard” in **Figure 4**) revealed the presence of Li ions in the RSF, which originated from the silk fibers’ dissolution in LiBr. The number of metal ions per single protein chain was calculated by dividing the measured weight of the detected ions by the total weight of the RSF sample and using the ions’ and fibroin’s molecular weight (see detailed calculation in Materials and Methods). We found that the number of Li ions per single fibroin protein chain is ∼60. It has been previously suggested that high concentrations of Li, which are not naturally present in the silk feedstock, prevent fibroin aggregation^10^.

Doping the “Standard” RSF solution with K or Na ions led to the replacement of Li ions with the dopant element. Interestingly, the total number of ions per fibroin chain remained relatively constant and only slight increased after the metal-doping procedure had been conducted. Each fibroin molecule was found to bind a total of 80–86 metal ions, indicative of the number of binding sites in each chain. These numbers roughly suit the ∼78 glutamic and aspartic acids in the fibroin sequence, mostly known to interact with metal ions. However, we found that doping RSF with Ca decreased the total number of ions in silk, reflecting the release of Li ions without their replacement by the same number of Ca ions. After Ca doping, the total number of binding ions per fibroin chain was measured to be 6.5–8, which is 10–12-fold lower than the monovalent dopant metal ions (i.e., K and Na). Thus, we can assume that the Ca ions have a much higher coordination number (10– 12 times higher) than monovalent ions when interacting with fibroin. Ca ions can form complexes with irregular geometry ligands and occupy more binding sites per fibroin chain, which might restrict the motion of the protein and prevent the formation of new conformations.

In agreement with other works^12,29^, our results (**Figure 3**) set the ratio of K and Ca ions per fibroin molecule in native silk feedstock to range between 10–50 and 1–40, respectively. To examine how different metal ion concentrations affect the silk feedstock’s flow properties, we doped the RSF with increasing K or Ca concentrations so as to generate protein-to-ion ratios of 1:3, 1:30, and 1:300.

The viscosity and viscoelasticity of the doped solutions were tested before and after two shear- and-relaxation cycles (**Figures 4 and supplementary Figures S7 and S8**). We observed that the initial pre-shear viscosity of RSF solutions decreases significantly after doping with either K or Ca (**Figures 4b and c**). This is similar to the finding of Laity et al. of an inverse correlation between K and Ca concentrations and the viscosity of silk feedstock^12^. The viscosity of non-doped RSF at the initial low shear rate of 1s^-1^ (**Figure 4d**) changes over time, starting from lower values that quickly increase and stabilize due to the applied shear. Without metal ions, which can increase the protein’s solubility^21^, the fibroin-soluble form destabilizes and tends to align and aggregate even at low shear rates. In contrast, the viscosity of metal-doped RSF does not significantly increase over time at low shear rates (**Figures 4e and f**).

Additionally, the initial (pre-shear) resistance of RSF to oscillatory deformation and the viscoelasticity values of the loss (G’) and storage (G’) modulus decrease as the ion’s (i.e., Ca or K) concentration increases (**supplementary Figures S7a-c, and e**). The ions’ concentrations also correlate with the RSF behavior as a fluid (as measured by way of Tan(delta) tests) (**supplementary Figure S7g**). Namely, higher ion concentrations result in a more liquid-like behavior of the fluid, as opposed to its solid-elastic behavior.

The shear flow was applied in two cycles of a gradual increase and decrease in shear rate, as seen in **Figures 4g-i**; the averaged dynamic viscosity (at a shear rate of 1s^-1^) after each step is presented in **Figures 4b and c**. Right after the *first* applied shear, the RSF viscosity increases significantly in all doped samples, but to a lesser extent in the RSF doped with higher Ca concentrations. When it is given time to relax at a low shear rate, non-doped RSF maintains its new higher viscosity, whereas K-doped RSF’s viscosity only slightly decreases (**Figure 4b**). However, K-doped RSF completely reverts back to its initial low viscosity when doped with high concentrations of Ca (1:30, 1:300, **Figure 4c**). Following the *second* applied shear, the viscosity increases once again to its previous high values in all the solutions. After the second relaxation time, the non-doped RSF and all K-doped RSFs maintain their high viscosities. The RSFs doped with high Ca concentrations (1:30, 1:300) revert to lower viscosities, but to ones higher than the initial values. The final “relaxed” viscosity remains fairly constant over time in all the RSF samples (**Figures 4d-f**).

The metal ions and shear similarly affect RSF viscoelastic properties (**supplementary Figure S7**). After applying shear, the overall resistance to deformation increases in all the samples, namely, both G’ and G’ rise (**supplementary Figures S7a**, **b**, **d, and f**). In addition, the storage modulus (G’) more significantly increases, and the material behaves more as a solid (as seen by the increase in Tan(delta), **supplementary Figures S7a**, **b, and h**), indicating that the protein has undergone fibrillation. However, as the Ca concentration increases (**supplementary Figures S7b**, **d**, **f, and h**), the RSF is more easily deformed (G’ and G’ decrease) and acts more as a liquid (Tan(delta) is greater than in other samples), even after high shear.

This partially reversible process of changes in RSF viscosity due to applied shear results from two simultaneous events: a molecular alignment during the high-acting shear that reverts back to a less ordered material after relaxation, and the shear-sensitive protein’s fibrillation that forms stable nano-fibrils. The different effects and events that occur during silk processing are best seen when the viscosity is plotted against the changing shear rates (**Figures 4g-i**). Initially, all the RSF samples behave similarly as a “shear thinning” fluid; their viscosity constantly decreases as the shear rate increases due to the disentangling and aligning of the molecular chains during the applied shear flow. However, at a critical shear rate (different for each dopant), the viscosity sharply rises (“step”) and assumes upon its stabilization a new decreasing trend; this “step” is a result of the silk protein’s shear sensitivity and is also affected by metal ions (**supplementary Figure S8**). In this sense, RSF solutions doped with increasing concentrations of Ca are much less sensitive to shear, and their critical shear rates are higher.

At high shear rates, the aligned and untangled molecular chains come into closer proximity until reaching a point where they interact and bind to each other, causing them to fibrillate and increase their bulk viscosity. For non-doped and K-doped RSF solutions, most proteins are irreversibly fibrillated after the first shear-relax cycle, as they mostly remain highly viscous even after relaxation (**Figures 4g and h**). However, Ca-doped RSF (**Figure 4i**) depicts a unique behavior: while the shear rate decreases after the *first* cycle, in the viscosity “steps” it reaches a lower value and ultimately returns to its original “pre-shear” values. During the *second* cycle, the same trend partially repeats: a “step” up at a high critical shear rate and a “step” down after lowering the shear rate. The interaction of Ca ions with the proteins prevents alignment of the chains, necessitating higher shear rates to achieve chain alignment, and consequently also inhibits the formation of nano-fibrils, allowing the protein to stay in its soluble form.

The overall effect of metal ions on RSF rheological characteristics suggests the following roles for metal ions during natural silk processing: In the silk gland’s middle parts, the metal ions appear at relatively high concentrations (**Figure 3**), thus reducing the silk feedstock viscosity. The interaction of fibroin with the metal ions, especially Ca, stabilizes its soluble form, making it less sensitive to shear and more suitable for storage. Once the feedstock reaches the anterior part, the ion concentrations fall sharply (**Figure 4**), causing an increase in the fluid’s viscosity and making it much more shear-sensitive. In this state, it is more prone to molecular alignment and fibrillation as it flows, two essential stages in the process of silk fiber formation.

### Metal ions modulate the mechanical properties of silk fibers

Spun silk fiber owes its non-standard ability to combine high stiffness, extensibility, strength and toughness to its hierarchical structure and non-covalent molecular interactions (mostly its hydrogen-bonded network). Previous studies have indicated that changes in the silk feedstock’s composition of metal ions, specifically in Ca and K, can affect the mechanics of the spun fibers^24–26,37^. However, whether the effect of metal ions on the fiber’s characteristics is further extended to the already spun fibers is not clear.

To elucidate this point, we doped degummed silk fibers with K or Ca (see Materials and Methods) and then examined their mechanical performance by tensile tests (**Figure 5**). The control treatment with distilled water slightly improved the mechanical properties of the fibers, indicating that the procedure did not harm the fibers; it kept the Young’s modulus and strain-to-failure at ∼10 Gpa and 18–19%, respectively, (**Figures 5b and c**) and increased the strength from ∼380 Mpa to ∼440 Mpa and the toughness from ∼52 MJ/m^3^ to ∼62 MJ/m^3^ (**Figures 5d and e**). In contrast, Ca doping significantly altered the fiber’s mechanical performance: it reduced Young’s modulus (∼8.5 Gpa), strength (∼268 Mpa), strain-to-failure (∼11.5%), and toughness (∼23 MJ/m^3^). Treatment with K had no significant effect on the strength (360 Mpa) and Young’s modulus (∼10 Gpa), but reduced the strain-to-failure (∼14%) and, consequently, the toughness (∼35 MJ/m^3^) of the fibers.

**Figure 5.**
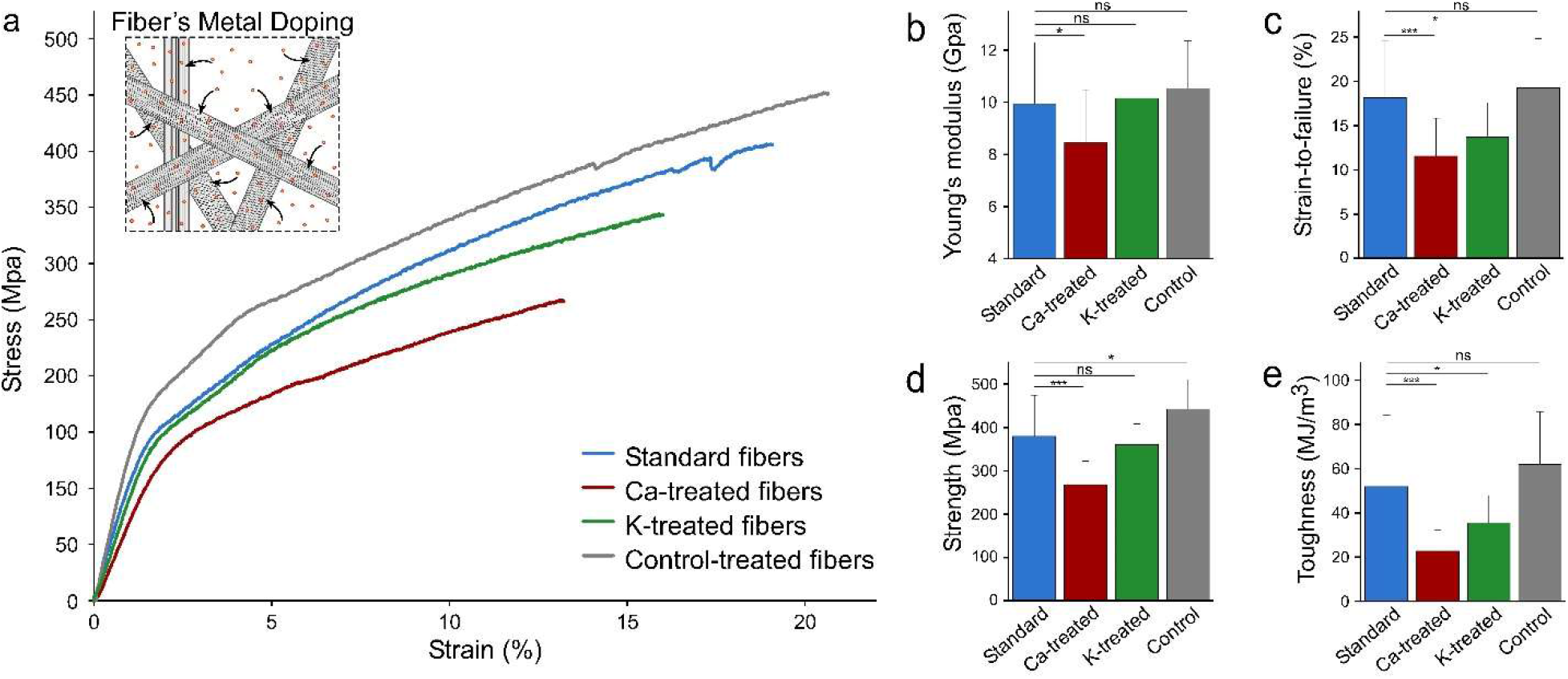
Mechanical properties of silk fibers doped with metal ions. *(a) Typical stress-strain curve of silk fibers before (standard) and after treatment with Ca or K metal ion or control (distilled water) solutions. Inset illustrates the doping of silk fibers with metal ions. (b-e) Column charts comparing the Young’s modulus (b), strain-to-failure (c), strength (d) and toughness (e) of the fibers subjected to the different treatments (standard and Ca, K and control treated). ns (not significant) - P>0.05, *-P<0.05, **-P<0.01, ***-P<0.001*.

In the case of fibroin, Ca at high concentrations acts as a chaotropic ion, destabilizing the protein’s solid β-sheet-rich form and increasing its solubilized random form^21,23^. Thus, it makes sense that the Ca treatment will alter the silk fiber’s structure and damage its mechanical properties, as has also been demonstrated by Wang et al.^25^.

As to K, it has been suggested that at mild concentrations, K serves as a cosmotropic agent, inducing fibroin fibrillation and aggregation^19,29^. In the context of the whole cocoon, including fibroin and sericin, the concentration of K has been shown to correlate with improved mechanical properties^25,29^. However, not only did the K treatment not improve the degummed fiber’s mechanical properties, it lowered the strain-to-failure (and thus the toughness). As has already been discussed hereabove, there is a logical inverse correlation between K’s concentration and its effect on silk protein’s mechanical properties during silk protein fibrillation (**Figure 4**) and on the fibroin core of the spun silk fiber (**Figure 2**). Thus, high concentrations of K interacting with fibroin fibers damage its mechanical properties.

Taken together, our results show that metal ions can affect silk fiber’s mechanical properties even after the completion of the spinning process.

### Conclusions

In this study, we analyzed metal ions’ composition and spatial distribution in silk fibers and *in-vivo* feedstock and examined their effect on silk fiber’s physical characteristics. Our results show that silk fibroin in the posterior and middle silk gland sections is exposed to high concentrations of metal ions, ensuring stable feedstock storage by reducing its viscosity and sensitivity to shear forces. By doing so, the ions (especially Ca) inhibit fibroin fibrillation and maintain its soluble form. Once the silk feedstock enters the gland’s anterior section, the metal ions’ concentrations in the silk feedstock fall dramatically, making fibroin more sensitive to shear forces. During its flow in the anterior section, the fibroin proteins are easily disentangled and aligned under the applied shear forces, increasing the feedstock viscosity, a change that causes the proteins to interact and start to fibrillate. The final silk fibers have a metal-deficient fibroin core and a sericin coating rich in metal ions. The composition of metal ions in the silk feedstock during spinning, and even in the already spun fibers, significantly modulates their mechanical performance.

The ions’ concentration and interaction with fibroin are carefully tuned during the process of natural silk production. They are essential for ensuring the material’s optimal properties for feedstock storage inside the silk gland, its liquid-to-solid transition and fibrillation, and the function of the final silk fibers. Our findings highlight the importance of metal ions in the material evolution of silk fibers and contribute to developing new concepts for modulating the properties of silk protein-based materials and other biomaterial designs.

## Acknowledgments

We thank Prof. Cedric Dicko from Lund University (Sweden), Prof. Gilad Haran and Prof. Jacob Klein for useful discussions and for comments on the manuscript. US acknowledges financial support from the Nella and Leon Benoziyo Center for Neurological Diseases. M.E.M. thanks Sergio Lombroso Fellowship (for Cancer Research) for financial support. In addition, US thanks the Perlman family for funding the Shimanovich Lab at the Weizmann Institute of Science: “This research was made possible in part by the generosity of the Harold Perlman Family”. The authors would like to acknowledge partial support from the GMJ Schmidt Minerva Center of Supramolecular Architectures at the Weizmann Institute, Mondry Family Fund for University of Michigan/Weizmann collaboration, Gerald Schwartz and Heather Reisman Foundation, WIS Sustainability and Energy Research Initiative (SAERI). This research was supported by a research grant from the Tom and Mary Beck Center for Advanced and Intelligent Materials at the Weizmann Institute of Science, Rehovot, Israel. The authors would like to thank Ofra Golani from the Department of Life Sciences Core Facilities, Weizmann Institute of Science, for planning and writing the ImageJ macro script to measure the thickness of the fibers for the mechanical tests. The authors would like to acknowledge the Scanning Electron Microscopy Unit at the Weizmann Institute of Science. The authors thank Ishai Dror and Stephen Fox from the Weizmann Institute of Science for performing the ICP-MS measurements. The authors are grateful to Natalie Page for English editing the manuscript.

## Author contributions

O.B. devised, designed, performed and analyzed the data of the experiments on metal ions quantifications, ICP-MS, SEM-EDS, Cryo-SEM-EDS, Rheology, tensile tests, silkworm dissection, and SDS-PAGE gels. O.B. drafted the manuscript and figures. E.S. supervised, guided and helped in performing all the SEM, Cryo-SEM and Cryogenic samples preparations, and the related results interpretation. D.E. helped in designing and performing the rheology, tensile tests and SDS-PAGE gels experiments and in data interpretation. I.K.A supervised and guided in performing the SEM and SEM-EDS imaging and in EDS data analysis and interpretation. I.C. performed and analyzed the data of the experiments on the metal ions binding to RSF and CD.

U.S. devised and supervised the project, contributed to the design and implementation of the research, to the analysis and interpretation of the results, and to the writing of the manuscript.

