## Supplementary data for "Metal Ions Guide the Production of Silkworm Fibers"

### Supplementary Information

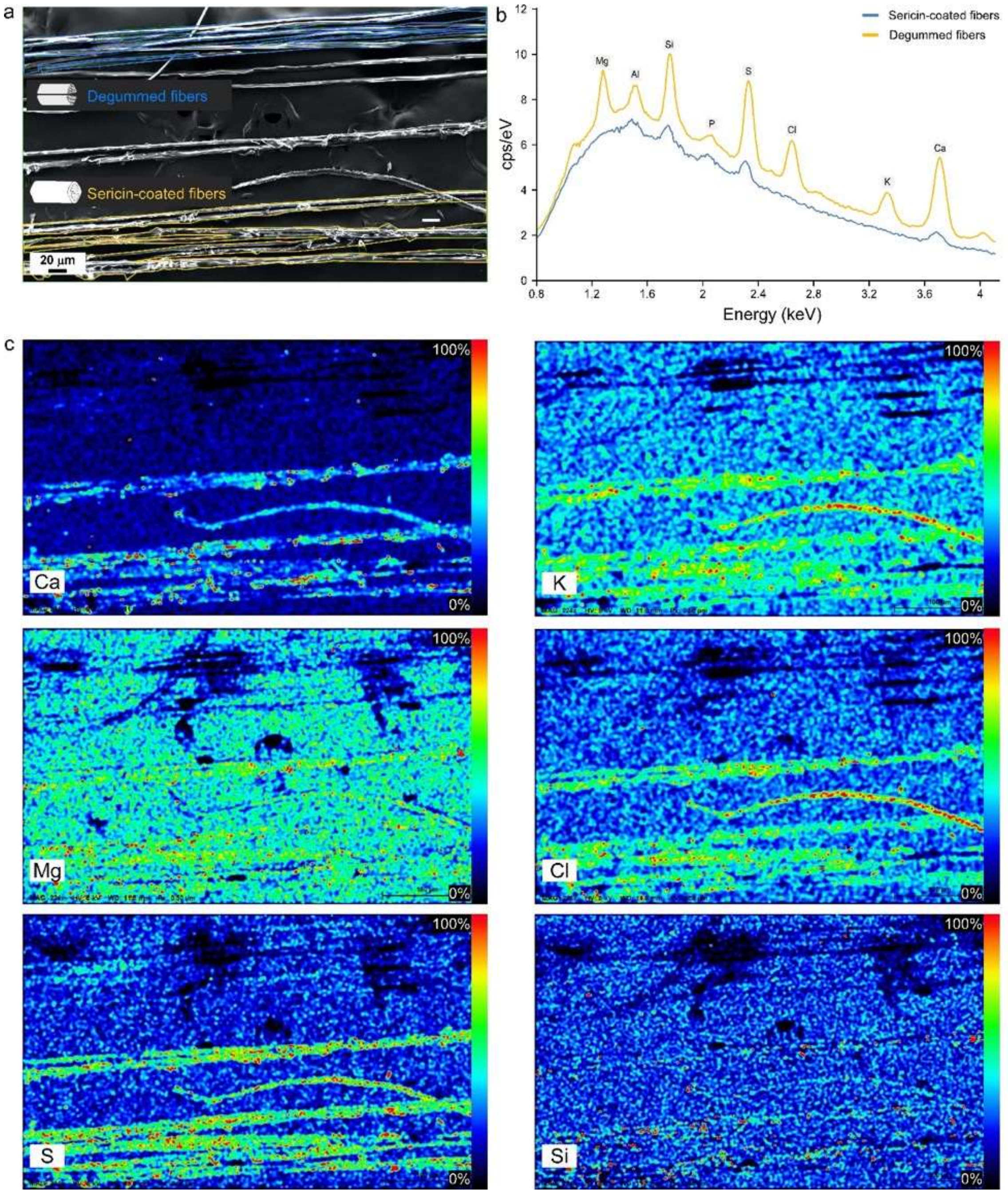


***Supplementary Figure S1: Representative images and SEM-EDS analysis of native (sericin-coated) and degummed silk fibers.*** *(a) Fiber regions before (orange) and after (blue) removing the sericin coating by degumming; scale bar 20 µm. (b) Characteristic X-ray spectrum of selected elements (Mg, Al, Si, P, S, Cl, K, Ca) in native and degummed silk fibers. (c) Normalized SEM-EDS intensity maps of the Ca, Mg, S, K, Cl and Si elemental distribution in the fibers. The EDS was obtained at an accelerating voltage of 8 kV.*

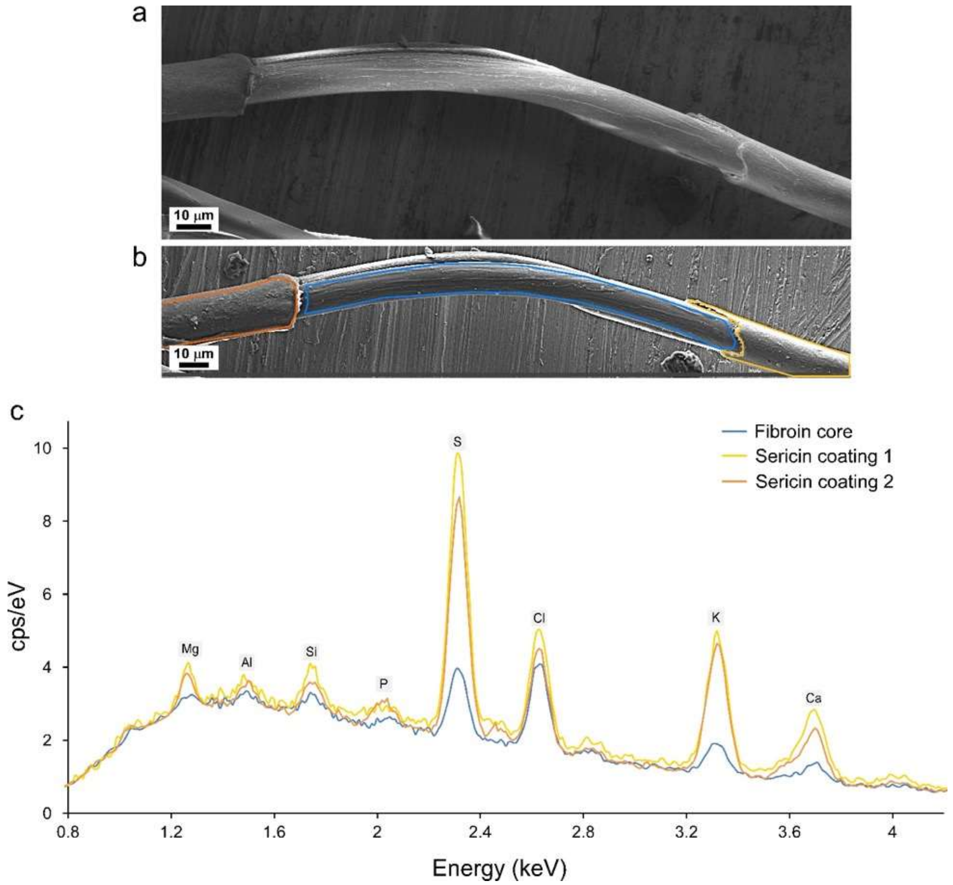


***Supplementary Figure S2 : Metal ions' distribution in silk fiber.*** *(a) SEM image of a silk fiber with a defective sericin coating; scale bar 10 μm. (b) Regions of interest in the fiber in (a): sericin-coated areas (yellow and orange), exposed uncoated fibroin (blue) and titanium surface (grey). (c) Characteristic elemental X-ray spectrum (EDS) in the regions of the silk fiber marked in (b). The EDS was obtained at an accelerating voltage of 8 kV.*

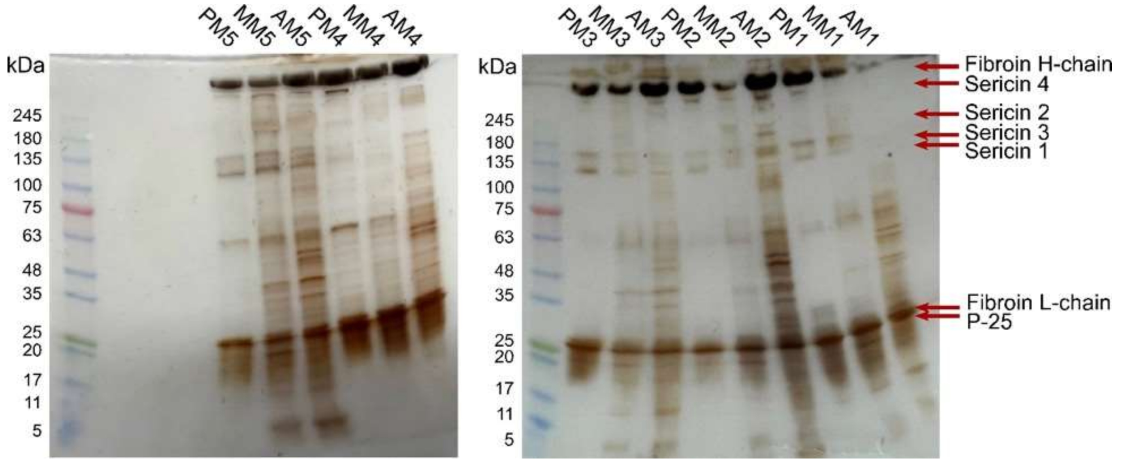


***Supplementary Figure S3:*** *SDS-gels of lumen protein extractions from Bombyx mori silk glands. Silk gland sections: AM - anterior middle, MM - middle middle, PM - posterior middle.*

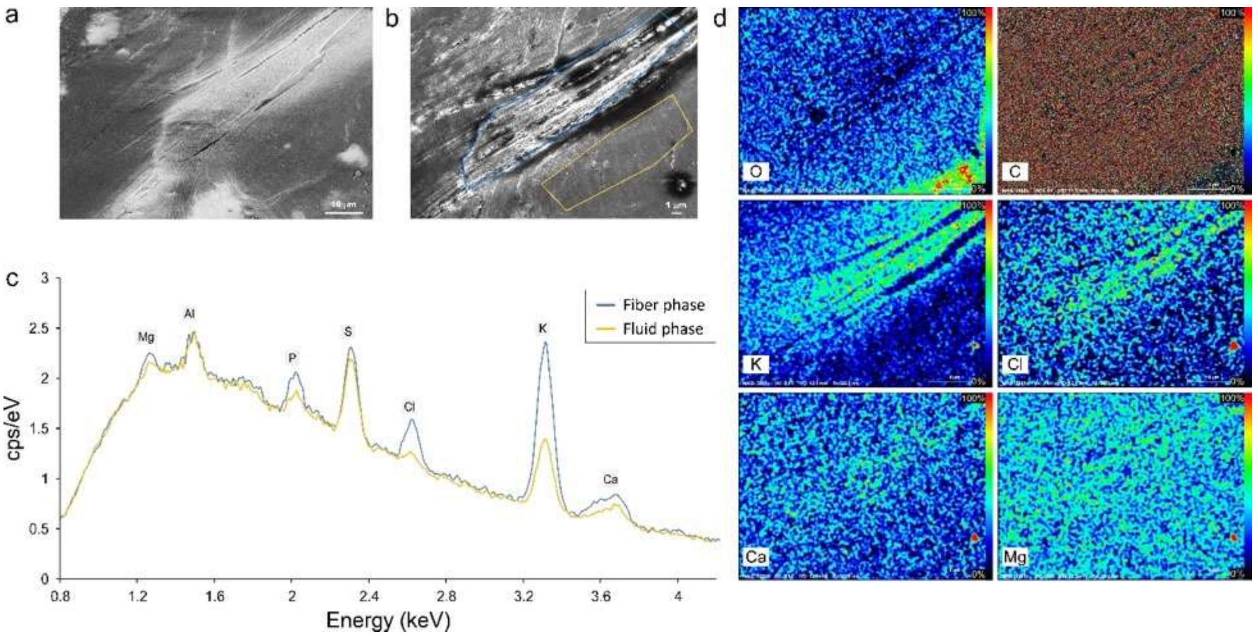


***Supplementary Figure S4: Distribution of ions in the front of the fiber-like structure formed in the silk gland, following high shear.*** *(a) SEM Image of a fiber-like structure formed in the silk gland following high shear; scale bar 10 µm. (b) Regions of interest in (a): the fiber-like phase (blue) and another lumen region (orange); scale bar 1 µm. (c) Characteristic elemental X-ray spectrum (EDS) in the fluid (orange) and fiber (blue) phases of the silk feedstock in the lumen. (d) Normalized SEM-EDS intensity maps of O, K, Ca, C, Cl, and Mg elemental distribution. The SEM-EDS was obtained with an accelerating voltage of 8 kV.*

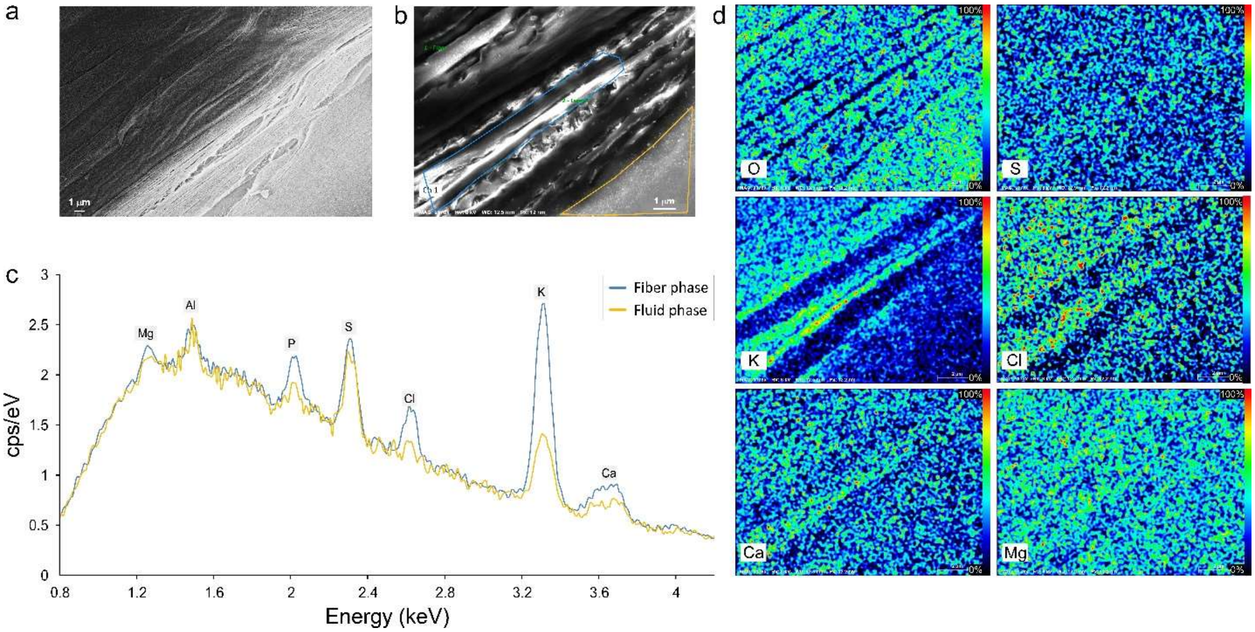


***Supplementary Figure S5: Ion distribution in the back of the fiber-like structure formed in the silk gland, following high shear.*** *(a) SEM image of a fiber-like structure formed in the silk gland following high shear; scale bar 1 µm. (b) Regions of interest: fiber-like phase (blue) and another lumen region (yellow). (c) Characteristic elemental X-ray spectrum (EDS) in the fluid (orange) and fiber (blue) phases of silk feedstock in the lumen. (d) Normalized SEM-EDS intensity maps of O, K, Ca, C, Cl, and Mg elemental distribution. The EDS was obtained with an accelerating voltage of 8 kV.*

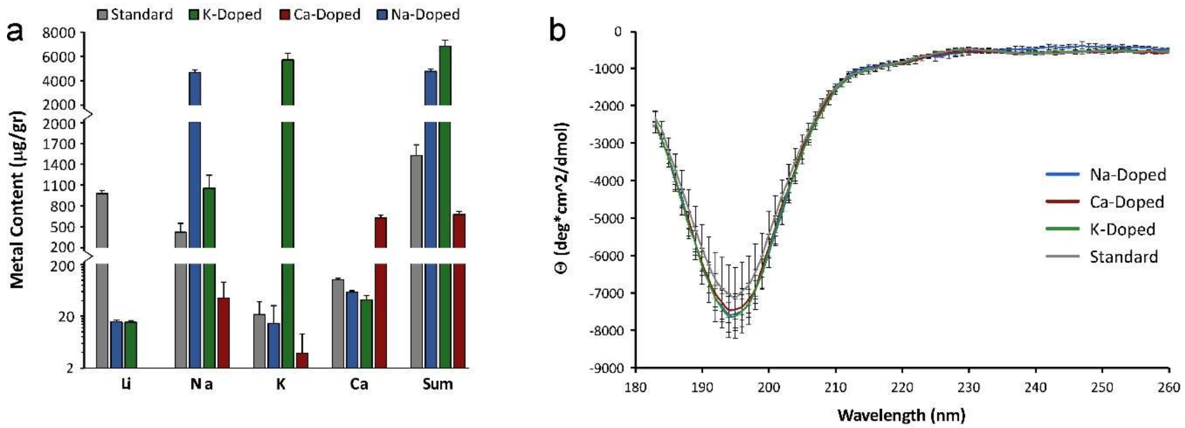


***Supplementary Figure S6: RSF dopping with the metal ions K, Ca and Na.*** *(a) ICP-MS elemental analysis of the RSF solutions before and after metal-doping or control (distilled water). (b) Circular dichroism (CD) spectrums of RSF after doping with Na, Ca, K or control (distilled water).*

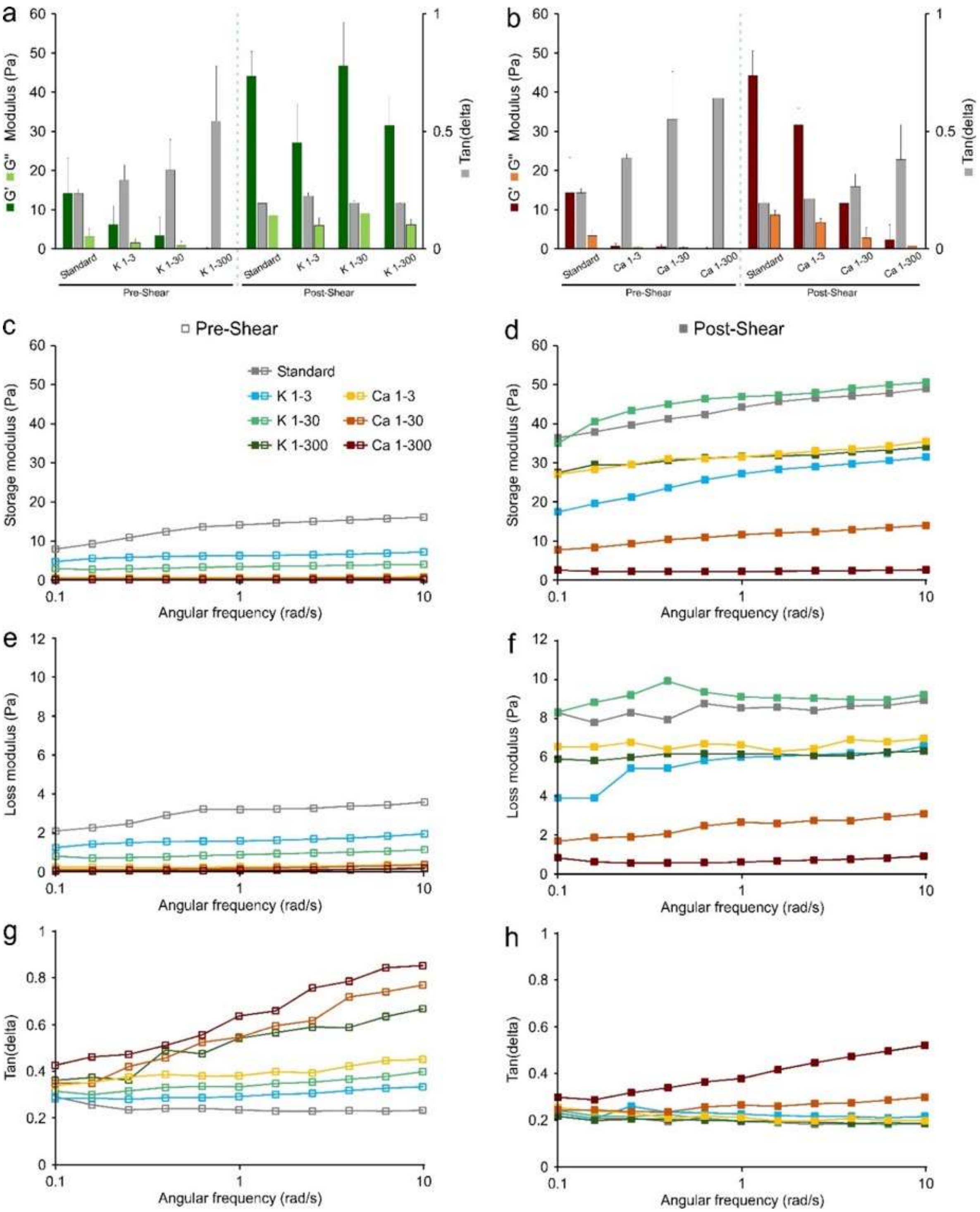


***Supplementary Figure S7: Viscoelasticity of RSF doped with K or Ca ions.*** *The averaged storage modulus (G''), loss modulus (G'') and Tan(delta) of K-doped (a) and Ca-doped (b) RSF samples before (pre) and after (post) applying high shear. (c-h) Oscillatory frequency sweep for metal-doped RSF presenting the change in G' (c,d), G'' (e,f) and Tan(delta) (g,h) before and after applying high shear.*

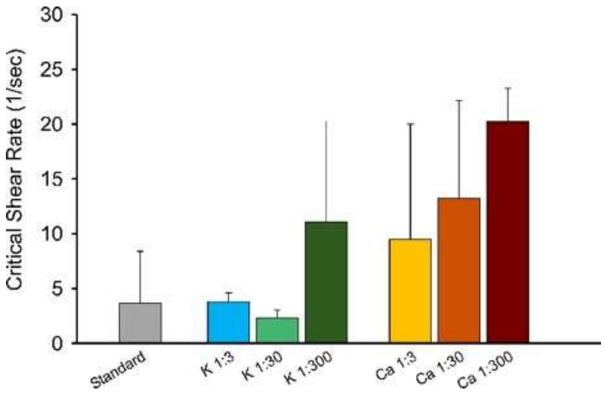


***Supplementary Figure S8:*** *The critical shear rate in which the trend in viscosity (as a function of the shear rate) shifts due to the shear sensitivity of fibroin. Different samples are RSF-doped with increasing concentrations of K or Ca, presented as the ratio between the protein molecules and ions.*
